# Kinase independent function of PI3Kγ modulates calcium re-uptake by regulating phospholamban function

**DOI:** 10.1101/2021.05.11.443650

**Authors:** Maradumane L. Mohan, Connner P. Witherow, Robert S. Papay, Yu Sun, Kate Stenson, Sathyamangla V. Naga Prasad

## Abstract

**Rationale:** Genetic deletion of Phosphoinositide 3-kinase (PI3Kγ) in mice (PI3Kγ^−/−^) results in increased cAMP levels and enhanced ventricular rate/contractility. Whether PI3Kγ plays a role in cardiac contractility by altering intracellular calcium recycling is not known.

**Objective:** To understand the mechanism of PI3Kγ mediated regulation of cardiac contractility.

**Methods and Results:** Caffeine treatment of adult cardiomyocytes from PI3Kγ^−/−^ mice showed significantly reduced calcium reuptake by sarcoendoplasmic reticulum (SR) indicating that PI3Kγ locally regulates SR function. This resulted in elevated levels of intracellular calcium for prolonged period following caffeine. Our findings show that delayed re-uptake of calcium was caused by changes in phosphorylation of phospholamban (PLN), a major regulator of SR calcium reuptake. PI3Kγ^−/−^ cardiomyocytes show significantly reduced PLN phosphorylation due to increase in PLN-associated protein phosphatase (PP) activity as reflected by decreased demethylated-PP2A. Consistently, the altered calcium regulation in the cardiomyocytes of PI3Kγ^−/−^ can be restored by inhibition of PP by okadaic acid. Unexpectedly, cardiomyocyate-specific overexpression of kinase-dead PI3Kγ PI3Kγ_inact_) in the global PI3Kγ^−/−^ cardiomyocytes normalized caffeine induced calcium reuptake, restored PLN phosphorylation, and decreased PLN-associated PP activity reflected by increased demethylated-PP2A.

**Conclusions:** These studies bring-to-fore an unrecognized regulation of PLN by PI3Kγ through PP2A with implications in deleterious cardiac remodeling as PI3Kγ is significantly upregulated following cardiac stress.

## INTRODUCTION

Calcium (Ca^2+^) is a divalent cation and its mobilization is universally important in regulation of cell function and signaling (Berridge, et al. 2003; Berridge, et al. 2000; Clapham 2007). It regulates plethora of cellular functions including muscle contraction, gene expression, hormone secretion, activation of kinases and phosphastases (Berridge, et al. 2003; Berridge, et al. 2000; Clapham 2007; Sheng, et al. 1990; West, et al. 2001; Wollheim and Sharp 1981). Ca^2+^ is the central regulator of excitation-contraction (EC) coupling in the heart (Berridge 2003; Bers 2002; Fabiato and Fabiato 1979; Fearnley, et al. 2011; Lederer, et al. 1990). EC coupling is activated by a relatively small Ca^2+^ influx from extracellular space triggering a quantitatively larger intracellular Ca^2+^ release from sarcoendoplasmic reticulum (SR) through ryanodine receptor (RyR2) channels (Bers 2004). This is followed by clearance of Ca^2+^ from cytosol by reuptake, majority of which is taken up by the SR through SR Ca2+ ATPase (SERCA2) pumps (Periasamy, et al. 2008; Periasamy and Huke 2001). The activity of SERCA2 is actively regulated by phospholamban (PLN), wherein unphosphorylated PLN binds and inhibits SERCA2 activity blocking the Ca^2+^ reuptake from cytoplasmic milieu. Phosphorylation of PLN results in its dissociation from SERCA2 leading to increased SR Ca^2+^ uptake (MacLennan, et al. 2003; MacLennan and Kranias 2003). Phosphorylation of PLN is mediated by cAMP-dependent protein kinase A (PKA) and Ca^2+^/Calmodulin-dependent protein kinase II (CaMKII), while, dephosphorylation of PLN is carried out by protein phosphatase 1 (PP1) and protein phosphatase 2A (PP2A) (MacLennan, et al. 2003; MacLennan and Kranias 2003) (Brittsan, et al. 2000; Brittsan and Kranias 2000). In this context, it is well known that methylation of PP2A leads to increased phosphatase activity, while demethylation results in reduction of activity (Stanevich, et al. 2011; Xing, et al. 2008). Dysregulation in these regulatory components underlies improper Ca^2+^ cycling leading to cardiac abnormalities including arrhythmias and heart failure (Bers, et al. 2003; Hasenfuss and Pieske 2002).

PI3Kγ belongs to family of lipid kinases that is activated downstream of G-protein coupled receptors (Stoyanov, et al. 1995; Vanhaesebroeck, et al. 2010) traditionally mediating Akt-dependent signaling critical for important cellular functions (Foster, et al. 2003; Rommel, et al. 2007; Ruckle, et al. 2006). Additionally, PI3Kγ also exhibits protein kinase activity (Czupalla, et al. 2003; Dhand, et al. 1994; Stoyanova, et al. 1997) (Naga Prasad, et al. 2005; Vasudevan, et al. 2011) and kinase-independent functions (Damilano, et al. 2010; Mohan, et al. 2013; Patrucco, et al. 2004). We have previously shown that PI3Kγ inhibits protein phosphatase activity via both kinase-dependent and kinase-independent mechanisms (Mohan, et al. 2017; Mohan, et al. 2013; Vasudevan, et al. 2011). In this context, the non-canonical scaffolding function of PI3Kγ assumes significance given the knowledge that PI3Kγ is expressed at very low levels in many organs including the heart (Ghigo and Li 2015; Martini, et al. 2014; Vanhaesebroeck, et al. 2010). Importantly, PI3Kγ is upregulated in conditions of stress/pathology like heart failure (Fougerat, et al. 2008; Patrucco, et al. 2004; Perino, et al. 2011; Perrino, et al. 2007) and cancer (Edling, et al. 2010; Xie, et al. 2013). Previous studies have shown that genetic deletion of PI3Kγ (PI3Kγ^−/−^) results in increased cardiac contractility (as measured by % FS) (Crackower, et al. 2002) and is associated with elevated levels of cAMP in the heart (Nienaber, et al. 2003). However, it is not known whether the changes in cardiac contractility is related to alterations in molecules regulating Ca^2+^ cycling. Given that cardiomyocyte contraction and EC coupling are tightly related to Ca^2+^ release from SR followed by reuptake, we tested whether PI3Kγ plays a role in regulating calcium cycling. Our comprehensive analysis of key players in Ca^2+^ cycling like RyR2, SERCA2 and PLN the shows that PLN is critically regulated by PI3Kγ that plays a pivotal role in releasing the inhibition on SERCA2. We show here that PI3Kγ releases the inhibition on SERCA2 by regulating PLN-associated phosphatases thereby, sustaining/maintaining PLN phosphorylation. Importantly, PI3Kγ engages kinase-independent mechanism instead of its kinase function to mediate inhibition of PLN-associated phosphatases. Intriguingly, this regulation seems to involve changes in the methylation state of PP2A. This shows unique and yet to be understood role of PI3Kγ in EC coupling that could have broader physiological impacts beyond its canonical role in regulation of GPCR function and downstream signaling.

## METHODS

### Animals

C57/Bl6 mice lacking PI3Kγ have been described previously (Nienaber et al 2003). Transgenic mice with cardiac-specific overexpression of kinase dead PI3Kγ mutant (PI3Kγ_inact_) (Nienaber et al 2003) in the PI3Kγ^−/−^ background (PI3Kγ_inact_/PI3Kγ^−/−^) have been described earlier (Mohan et al 2013). Experiments involving live animals were performed in accordance with institutional and national guidelines and regulations, as approved by Cleveland Clinic Institutional Animal Care and Use Committee.

### Isolation of cardiomyocytes

Adult mouse cardiomyocytes were isolated using previously described procedure (Li, D., Wu, J., Bai, Y., Zhao, X., Liu, L. 2014 J. Vis. Exp. (87), e51357). Briefly, the myocytes were separated in a high potassium buffer from the isolated heart by modified Langendorff perfusion (Buffer: 113 mM NaCl, 4.7 mM KCl, 0.6 mM KH_2_PO_4_, 0.6 mM Na_2_HPO_4_, 1.2 mM MgSO_4_, 10 mM Na-HEPES, 12 mM NaHCO_3_, 10 mM KHCO_3_, 0.032 mM phenol red, 30 mM taurine, 10 mM 2,3-Butanedione monoxime, and 5.5 mM glucose) and digestion with type II collagenase (300 U/mL in perfusion buffer containing 50 μM CaCl_2_). The myocytes were separated from other cell types by centrifugation.

### Measurement of intracellular calcium (Ca^2+^) transients

Intracellular Ca^2+^ transients were measured using the IonOptix simultaneous Contractility and Calcium system. The IonOptix Contractility and Calcium Assay protocol was adapted to best suit the experimental design and the type of cells that were being used for the study. Isolated mouse cardiomyocytes were incubated with Fura-2 AM (200 μM) for 15 min at 37 °C. Following incubation, the cells were washed twice for 5 minutes with Tyrodes solution (134 mM NaCl, 12 mM NaHCO_3_, 2.9 mM KCl, 0.34 mM Na_2_HPO_4_, 1 mM MgCl_2_, 10 mM HEPES pH 7.4, 1.2 mM CaCl_2_). The Fura-2-loaded cardiomyocytes were added to the perfusion chamber. The cardiomyocytes were paced at 3.0 Hz stimulation using the MyoPacer field stimulator (11.0 Volts). Using the 40X objective lens, healthy cardiomyocytes (rod shaped, well striated) were identified and used for recording contractility traces and Ca^2+^ transients. The cardiomyocytes were centered to the field of view and the background was minimized by adjusting the aperture. The Ca^2+^ recording was measured using the IonOptix system through the IonWizard software wherein the Calcium-Numeric-subtracted ratio of Fura-2 AM fluorescence (excitation at 360/380 nm and emission at 510 nm) was collected. A collection of similar transients for a period of 60 seconds was used for data analysis.

### Chemical Reagents

Caffeine was obtained from Sigma. Phospho-RyR2 (S2809) (1:1000), phospho– SERCA2 (S38) were obtained from Badrilla. RyR2 (1:1000), SERCA2 (1:1000), phospho-PLN (1:1000), and PLN (1:1000) antibodies were obtained from Cell Signaling Technology. PI3Kγ (1:1000) and PP1 (1:1000) antibodies were from Santacruz Biotechnology. PP2Ac (1:2000) antibody was obtained from Millipore. Okadaic Acid was obtained from Sigma.

### Wetern blotting and immunoprecipitation

Standard procedures for western immunoblotting and immunoprecipitations were followed. Protein extracts were prepared from hearts using either NP-40 lysis buffer (20 mM Tris pH 7.4, 137 mM NaCl, 1 mM PMSF, 20% Glycerol, 10 mM NaF, 1 % NP-40) for immunoblotting or TritonX100 lysis buffer (20 mM Tris pH 7.4, 300 mM NaCl, 1 mM PMSF, 20% Glycerol, 0.8 % TritonX 100) for immunoprecipitation experiment. Both the buffers contained protease and phosphatase inhibitor cocktails (Sigma). Lysates were cleared by centrifugation at 13,000 rpm for 15 min at 4°C. Supernatants were used for immunoblotting analysis or for immunoprecipitation with the indicated antibodies.

### SR fractionation

SR vesicles from hearts were isolated using differential centrifugation as previously described (Ref). Briefly, hearts were homogenized in phosphatase assay buffer (50 mM Tris-HCl, pH 7.0; 100 μM CaCl_2_) containing protease inhibitor cocktail. The homogenates were centrifuged at 14000 g for 20 min, supernatents were centriguged at 16000 g for 20 min and the resulting supernatents were centrifuged at 45000 g for 45 min to obtain SR vesicles as pellets. The SR pellets were resuspended in phosphatase assay buffer and utilized for phosphatase assay or immunoprecipitation.

### Phosphatase assay

Phosphatase activity was measured using phosphatase assay kit (Millipore). Immunoprecipitated samples were resuspended in phosphate-free assay buffer, and the assay was done according to the manufacturer’s protocol (Vasudevan et al 2011).

### Statistical Analysis

All data were expressed as mean±SEM (n≥3 experiments performed under identical conditions). ANOVA was used for multiple comparisons of the data. Statistical analyses were performed using GraphPad Prism© and the significance between the treatments were determined by Student *t-test*. A p-value less than < 0.05 was considered statistically significant.

## RESULTS

PI3Kγ plays a role in calcium reuptake by Sarcoendoplasmic Reticulum (SR) Previous studies have shown that PI3Kγ^−/−^ mice have higher ventricular contractility (Alloatti, et al. 2005; Crackower, et al. 2002; Kerfant, et al. 2005). In order to understand this phenomenon further, we assessed the contractility of cardiomyocytes isolated from WT and PI3Kγ^−/−^ mice. When compared to the WT the cardiomyocytes contractility was higher and the relaxation time was increased under basal conditions in PI3Kγ^−/−^ mice (**Fig 1A**). However, this phenomenon was recovered when the cardiomyocytes were stimulated with isoproterenol mediated adrenergic stimulation (**Fig S1**) suggesting in the myocytes of PI3Kγ^−/−^ mice. Since calcium cycling in the cardiac myocytes plays a key role in cardiomyocyte contractility, we assessed caffeine-induced calcium release/reuptake in isolated adult cardiomyocytes from WT and PI3Kγ^−/−^ mice using IonOptix system. Caffeine (10 mM) treatment resulted in robust calcium release followed immediately by effective reuptake in the WT adult cardiomyocyte (**Fig. 1B**). When compared to WT cardiomyocytes (**Fig 1B**), marked changes in the calcium cycling were observed in PI3Kγ^−/−^ cardiomyocytes (**Fig 1C**). The PI3Kγ^−/−^ cardiomyocytes were characterized by robust calcium release but lagged in calcium reuptake (**Fig 1C**) and interestingly, never reached back to the baseline. In order to further understand if the changes were in the release of calcium reserve or reuptake of the calcium by the SR, time taken to peak calcium level in the cytoplasm (a measure of calcium release) and time taken to baseline from peak calcium levels in the cytoplasm (a measure of calcium reuptake by SR) were calculated. Significant changes were not observed in average time to peak between WT and PI3Kγ^−/−^ myocytes (**Fig 1D**) suggesting that release of calcium from SR is not altered due to the absence of PI3Kγ. However, the average time to baseline was significantly increased in PI3Kγ^−/−^ mycotyes when compared to WT (**Fig 1E**). These data indicate that calcium reuptake by SR post release is significantly delayed in PI3Kγ^−/−^ mice leading to calcium overload in the myocytes.

**Fig 1.**
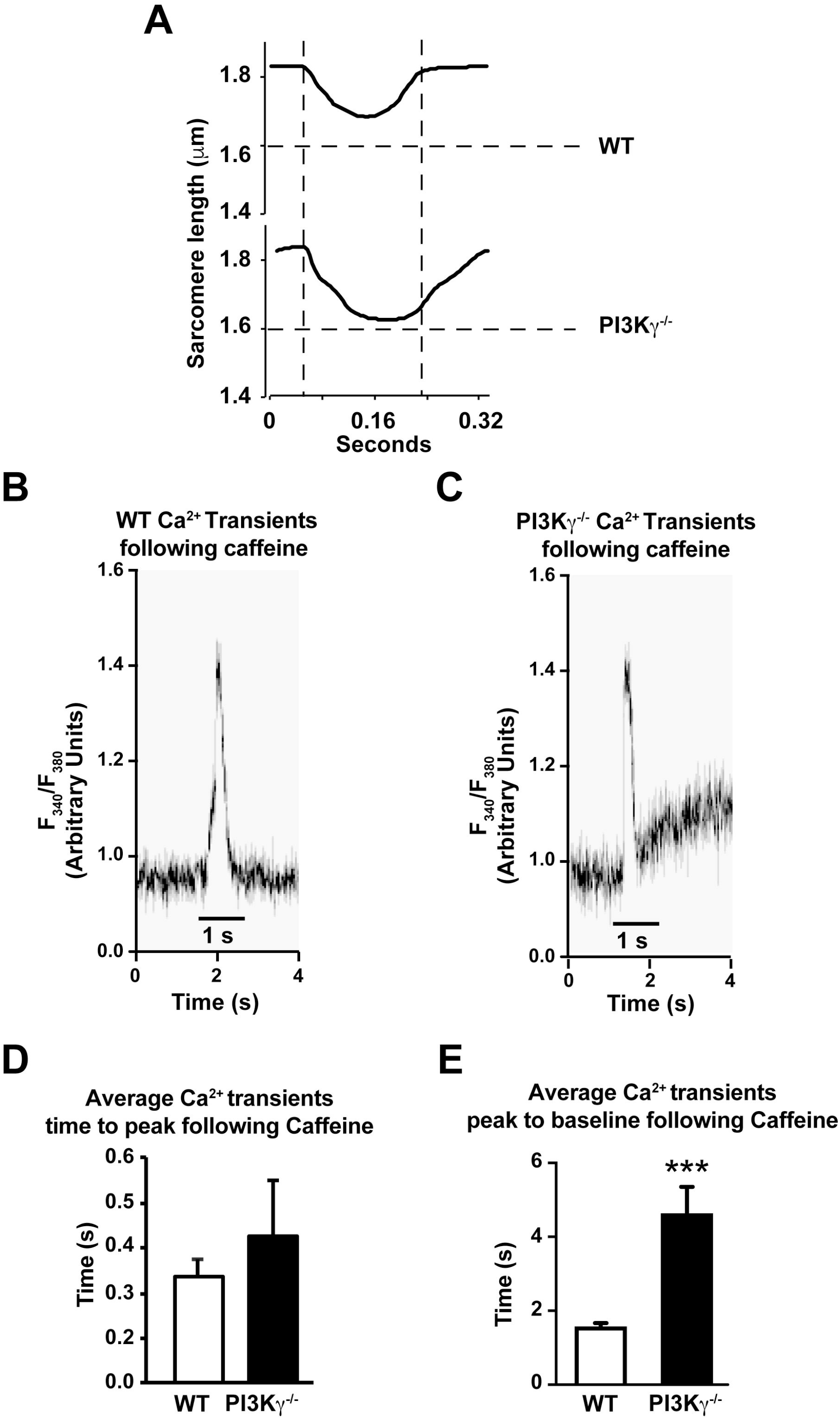
Caffeine induced calcium cycling is altered in PI3Kγ^−/−^ mice. **(A)** Average of cardiomyocyte contractility traces of WT and PI3Kγ^−/−^ cardiomyocytes (n=3 mice; 10-15 myocytes per mouse; 90-100 contractions/myocyte). **(B)** Representative image of calcium traces in cardiomyocytes isolated from WT mice analyzed for caffeine induced calcium cycling. **(C)** Representative image of calcium transients in cardiomyocytes isolated from PI3Kγ^−/−^ mice analyzed for caffeine induced calcium cycling. **(D**) Average calcium transients time to peak (measure of rate of calcium release from SR) for WT and PI3Kγ^−/−^ cardiomyocytes (n=3 mice; 10-15 myocytes/mouse; 180-200 contractions/myocyte). **(E)** Average calcium transients time to baseline from peak (measure of rate of reuptake of calcium by SR) for WT and PI3Kγ^−/−^ cardiomyocytes (n=3 mice; 10-15 myocytes/mouse; 180-200 contractions/myocyte). ***P < 0.001 compared to the WT.

### PI3Kγ regulates Phospholamban (PLN) phosphorylation

Given the intriguing observation of the increased intracellular calcium levels in the cytoplasm, we investigated the underlying molecular mechanisms contributing to this phenomenon. Based on the known regulators of calcium cycling, the delayed clearance of calclium from the cytoplasm in the PI3Kγ^−/−^ myocytes could be due to leaky Ryanodine receptors (RyR2) on SR or alterations in the function of SERCA2 pump. To test whether there is altered RyR2 function, we measured RyR2 phosphorylation levels by western immunoblotting. However, no differences in RyR2 phosphorylation were observed in the hearts of PI3Kγ^−/−^ mice compared to the WT indicating the normal functioning of RyR2 in these mice (**Fig 2A**). Similarly, immunoblotting for phosphorylation of SERCA2 showed no differences between PI3Kγ^−/−^ and WT mice suggesting that alterations in the SERCA2 itself is not the reason for delay in calcium reuptake (**Fig 2B**). SERCA2 function is known to be regulated by PLN, wherein unphosphorylated PLN binds to SERCA2 and inhibits SERCA2 function (MacLennan and Kranias 2003). Therefore, we probed the cardiac lysates for phospho-PLN levels. Surprisingly, significant loss of phosphorylation in PLN was observed in PI3Kγ^−/−^ when compared to WT (**Fig 2C & D**). These data show that PI3Kγ is required for PLN phosphorylation and thereby, normal functioning of SERCA2 to prevent dysregulation of calcium cycling in the myocytes.

**Fig 2.**
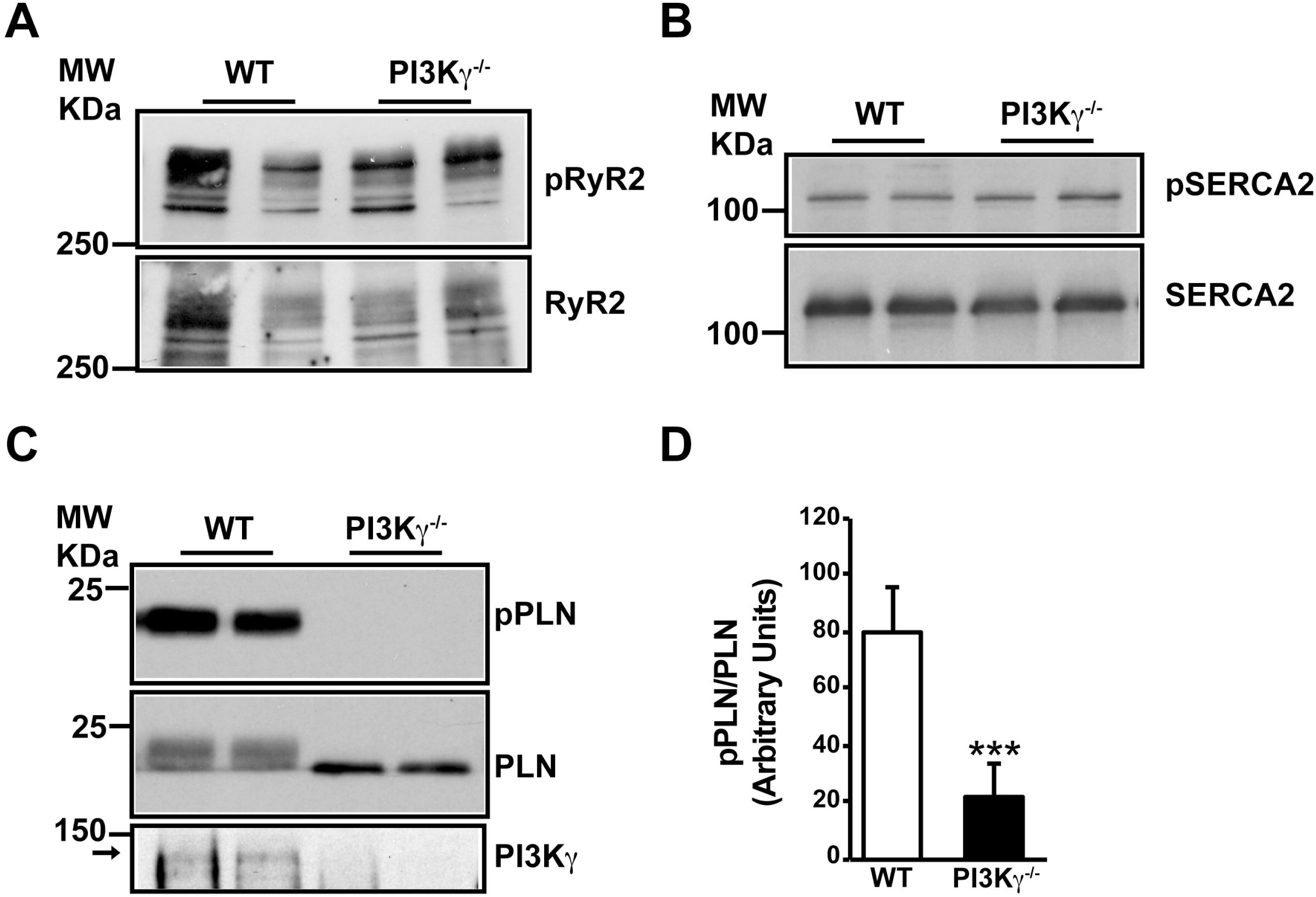
PI3Kγ dependent calcium cycling is through regulation of phospholamban phosphorylation. **(A)** Total cardiac lysates from wild-type (WT) and PI3Kγ^−/−^ mice were immunoblotted with anti–phospho-RyR2 antibody (pRyR2). The blots were stripped and reprobed for total RyR2 (lower panel). **(B)** Total cardiac lysates from WT and PI3Kγ^−/−^ mice were immunoblotted with anti–phospho-SERCA2 antibody (pSERCA2). The blots were stripped and reprobed for total SERCA2 (lower panel). **(C)** Total cardiac lysates from WT and PI3Kγ^−/−^ mice were immunoblotted with anti–phospho-PLN antibody (pPLN). The blots were stripped and reprobed for total PLN (lower panel). **(D)** Amalgamated densitometry of PLN phosphorylation is shown as bar graph. n = 6-8 mice, and each lane represents one mouse. ***P < 0.001 compared to the WT.

### PI3Kγ promotes PLN phosphorylation by inhibiting protein phosphatase

Since our previous studies have shown that PI3Kγ^−/−^ plays a critical role in regulating phosphatase function (Mohan, et al. 2017; Mohan, et al. 2013; Vasudevan, et al. 2011), we tested whether the loss of PLN phosphorylation was due to elevated phosphatase activity. SR fractions isolated from the hearts of WT and PI3Kγ^−/−^ mice were assessed for phosphatase activity using malachite green based phosphatase assay. The phosphatase activity on SR membrane was significantly increased in PI3Kγ^−/−^ mice when compared to WT mice (**Fig 3A**). We further tested if the change in phosphatase activity was specific to PLN by immunoprecipitating PLN from cardiac lysates and measuring PLN associated phosphatase activity. Similar to SR phosphatase activity, PLN associated phosphatase activity was significantly higher in PI3Kγ^−/−^ mice in comparison to WT mice (**Fig 3B**). These data indicate that PI3Kγ inhibits protein phosphatase at SR as its absence in the PI3Kγ^−/−^ leads significant increase in phosphatase activity. Thus, PI3Kγ is able to maintain calcium reuptake by SERCA2 through regulation of PLN phosphorylation uniquely through phosphatase mediated mechanisms.

**Fig 3.**
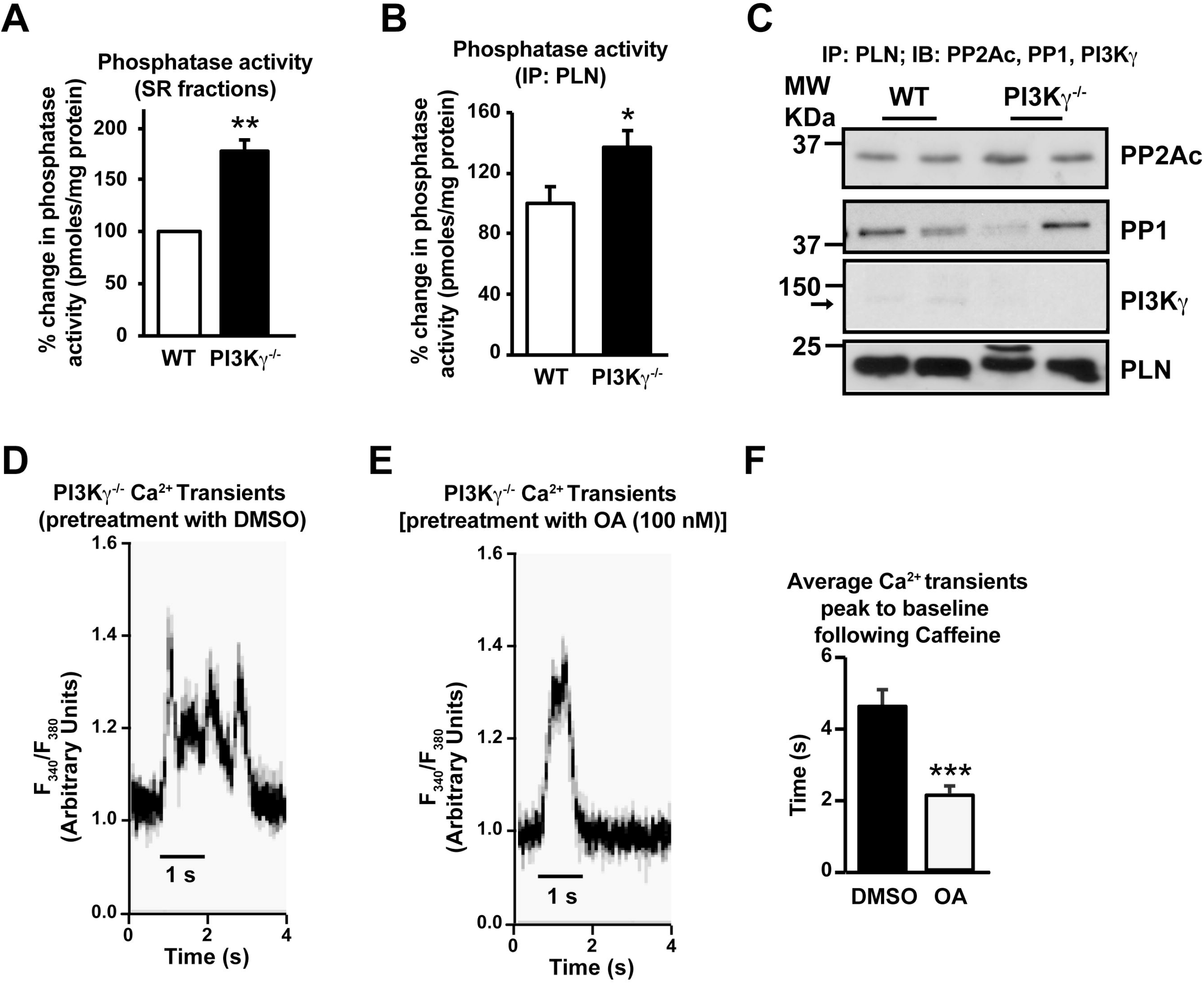
PI3Kγ controls calcium cycling through regulation of phosphatase-PLN axis. **(A)** In vitro phosphatase assays were carried out on SR fractions of WT and PI3Kγ^−/−^ mice hearts. Data presented as percent change of phosphtase activity in PI3Kγ^−/−^ over WT. n = 5 to 6 mice per genotype. **P < 0.01 compared to WT. **(B)** In vitro phosphatase assays were carried out on PLN immunoprecipitates from cardiac lysates of WT and PI3Kγ^−/−^ mice. Data presented as fold change in phosphatase activity in PI3Kγ^−/−^ over WT. n = 5 to 6 mice per genotype. *P < 0.05 compared to WT. **(C)** PLN immunoprecipitates from cardiac lysates of WT and PI3Kγ^−/−^ mice were immunoblotted for coimmunoprecipitating PP2Ac, PP1 and PI3Kγ and immunoblotted for PLN to show pull down of PLN by the antibody. **(D)** Representative image of calcium transients in cardiomyocytes isolated from PI3Kγ^−/−^ mice analyzed for caffeine induced calcium cycling following DMSO pretreatment. **(E)** Representative image of calcium transients in cardiomyocytes isolated from PI3Kγ^−/−^ mice analyzed for caffeine induced calcium cycling following OA pretreatment. **(F)** Average calcium transients time to baseline (measure of rate of reuptake of calcium by SR) for PI3Kγ^−/−^ cardiomyocytes following DMSO or OA pretreatment (n=3 mice; 10-15 myocytes/mouse; 180-200 contractions/myocyte). ***P<0.001 OA v/s DMSO.

In order to understand if the alterations in the phosphatase activity is due to changes in association of protein phosphatases with PLN, PLN was immunoprecipitated and immunoblotted for co-immunoprecipitating phosphatases (PP1 and PP2A). Interestingly, despite increase in PLN-associated phosphatase activity in the PI3Kγ^−/−^ hearts, there were no significant differences in the interaction of PP1 or PP2A with PLN in PI3Kγ^−/−^ hearts compared to the WT hearts (**Fig 3C**). Interestingly, PI3Kγ interaction was observed in WT heart which was absent in the PI3Kγ^−/−^ mice hearts (**Fig 3C)** suggesting a unique mechanism of PLN-associated protein phosphatase regulation.

To determine whether regulation of PLN phosphorylation through protein phosphatases by PI3Kγ underlies the altered calcium reuptake, adult PI3Kγ^−/−^ cardiomyocytes were pretreated with phosphatase inhibitor okadaic acid (OA). Following pretreatment, caffeine induced calcium release and reuptake were measured in PI3Kγ^−/−^ cardiomyocytes. Consistently, there was significant delay in the calcium reuptake in vehicle pretreated PI3Kγ^−/−^ cardiomyocytes (**Fig 3D**). This delay in calcium reuptake was significantly reversed following pretreatment of PI3Kγ^−/−^ cardiomyocytes with OA (**Fig 3E**). Corespondingly, the calcium reuptake average time to baseline was significantly reduced in cardiomyocytes following OA pretreatment (**Fig 3F**) suggesting that PI3Kγ regulation of PLN and calcium cycling is through inhibition of protein phosphatases.

### Calcium cycling regulation by PI3Kγ is kinase-independent

To test whether kinase activity of PI3Kγ is required for calcium cycling, we generated a unique mouse line with cardiomyocyte-specific overexpression of kinase-dead PI3Kγ mutant (PI3Kγ_inact_) in the PI3Kγ^−/−^ background (PI3Kγ_inact_/PI3Kγ^−/−^). Consistent with observations presented in Fig 1, PI3Kγ^−/−^ myocytes showed altered calcium cycling (**Fig 4B**) compared to WT myocytes (**Fig 4A**). However, unexpectedly the calcium cycling in the PI3Kγ_inact_/PI3Kγ^−/−^ cardiomyocytes was restored to a pattern (**Fig 4C**) similar to the WT cardiomyocytes (**Fig 4A**). No significant difference in average time for calcium to peak was observed in these cardiomyocytes (**Fig 4D**), but the average calcium reuptake time to baseline was selectively restored in PI3Kγ_inact_/PI3Kγ^−/−^ myocytes (**Fig 4E**). These findings show that the delayed reuptake of calcium observed in PI3Kγ^−/−^ could be rescued by kinase-independent function of PI3Kγ. To test whether kinase-independent function of PI3Kγ rescues PLN phosphorylation, cardiac lysates were immunoblotted for pPLN. Consistent with normalization of calcium reuptake, PLN phosphorylation was also restored by PI3Kγ_inact_ expression (**Fig 4F & G**) further supporting the kinaseindependent role of PI3Kγ in SR calcium cycling.

**Fig 4.**
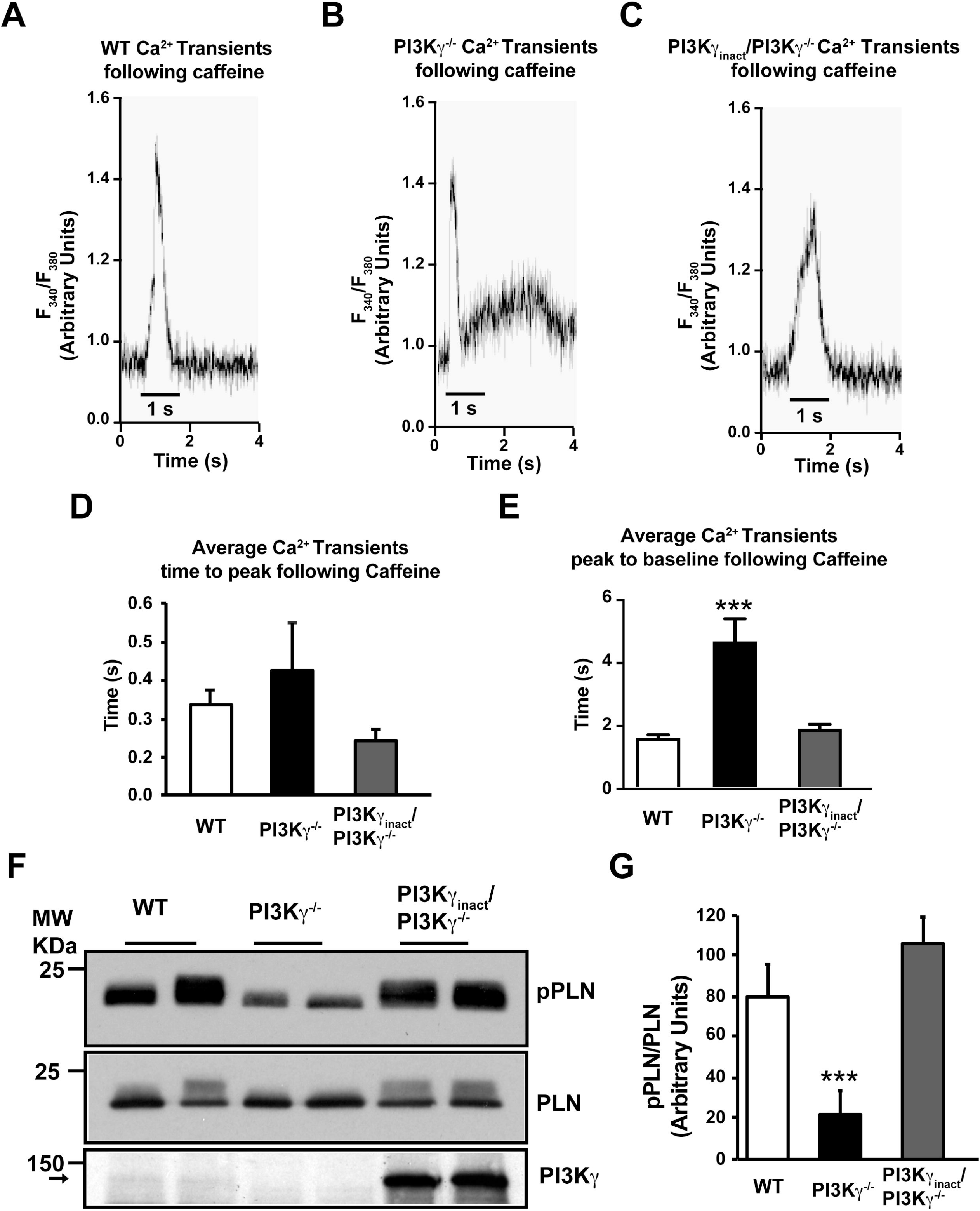
Calcium cycling is regulated by kinase-independent fumction of PI3Kγ. **(A)** Representative image of calcium transients in cardiomyocytes isolated from WT mice analyzed for caffeine induced calcium cycling. **(B)** Representative image of calcium transients in cardiomyocytes isolated from PI3Kγ^−/−^ mice analyzed for caffeine induced calcium cycling. **(C)** Representative image of calcium transients in cardiomyocytes isolated from PI3Kγ_inact_/PI3Kγ^−/−^ mice analyzed for caffeine induced calcium cycling. **(D)** Average calcium transients time to peak (measure of rate of calcium release from SR) for WT, PI3Kγ^−/−^ and PI3Kγ_inact_/PI3Kγ^−/−^ cardiomyocytes (n=3 mice; 10-15 myocytes/mouse; 180-200 contractions/myocyte). **(E)** Average calcium transients time to baseline from peak (measure of rate of reuptake of calcium by SR) for WT and PI3Kγ^−/−^ cardiomyocytes (n=3 mice; 10-15 myocytes/mouse; 180-200 contractions/myocyte). ***p<0.001 compared to WT and PI3Kγ_inact_/PI3Kγ^−/−^. **(F)** Total cardiac lysates from WT and PI3Kγ^−/−^ mice were immunoblotted with anti–phospho-PLN antibody (pPLN). The blots were stripped and reprobed for total PLN (lower panel). **(G)** Cumulative densitometric data for PLN phosphorylation is shown as bar graph. n = 6-8 mice, and each lane represents one mouse. ***P < 0.001 compared to the WT and PI3Kγ_inact_/PI3Kγ^−/−^.

### Regulation of PLN phosphatase activity by kinase-independent function of PI3Kγ controls assembly of SERCA2-PLN complex

To determine whether normalization of calcium cycling in the PI3Kγ_inact_/PI3Kγ^−/−^ is also associated with concomitant changes in the phosphatase activity, SR fractions were isolated and assessed for phosphatase activity. The phosphatase activity in the SR fraction of PI3Kγ_inact_/PI3Kγ^−/−^ was normalized to WT levels (**Fig 5A**) and similarly, PLNassociated phosphatase activity was also reduced in PI3Kγ_inact_/PI3Kγ^−/−^ mice compared to PI3Kγ^−/−^ mice (**Fig 5B**). Despite normalization of phosphatase activity in the PI3Kγ_inact_/PI3Kγ^−/−^, no significant differences in the co-immunoprecipitating PP1 or PP2A were observed upon PLN immunoprecipitation (**Fig 5C**). Since PI3Kγ inhibits PP2A activity kinase-independently by blocking PP2A methylation (Mohan, et al. 2013) we probed for demethylation levels of PP2A on SR fractions by western immunoblotting. We observed significantly lower abundance of dimethyl-PP2A in PI3Kγ^−/−^ mice in comparison to WT and PI3Kγ_inact_/PI3Kγ^−/−^ mice (**Fig 5D & E**). These data suggest higher level of PP2A methylation in PI3Kγ^−/−^ accounting for elevated phosphatase activity.

**Fig 5.**
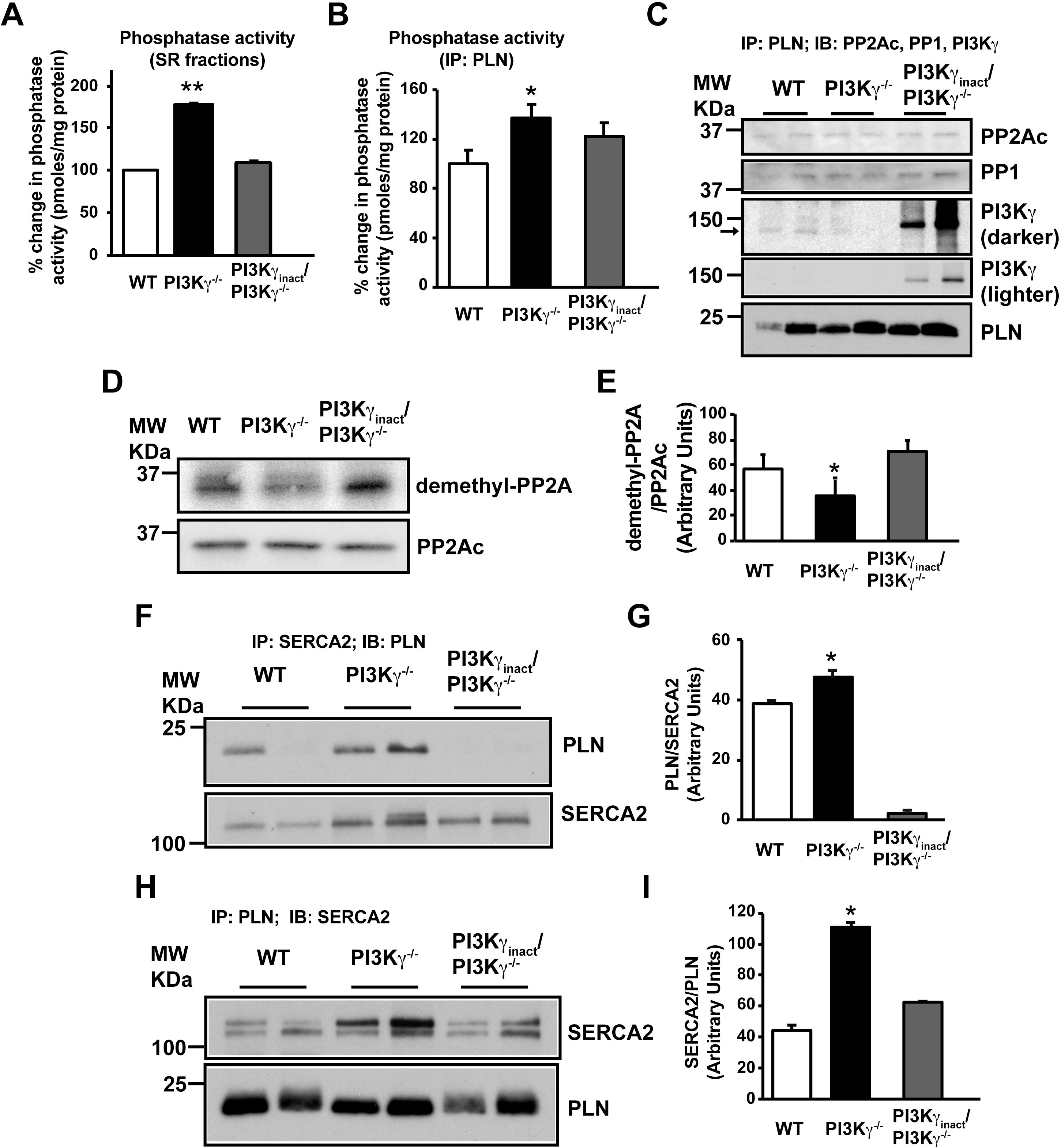
Kinase-independent function of PI3Kγ regulates phosphatase-PLN axis. **(A)** In vitro phosphatase assays were carried out on SR fractions of WT, PI3Kγ^−/−^ and PI3Kγ_inact_/PI3Kγ^−/−^ mice hearts. Data presented as percent change of phosphtase activity in PI3Kγ^−/−^ and PI3Kγ_inact_/PI3Kγ^−/−^ over WT. n = 5 to 6 mice per genotype. **P < 0.01 compared to WT and PI3Kγ_inact_/PI3Kγ^−/−^. **(B)** In vitro phosphatase assays were carried out on PLN immunoprecipitates from cardiac lysates of WT, PI3Kγ^−/−^ and PI3Kγ_inact_/PI3Kγ^−/−^ mice. Data presented as fold change in phosphatase activity in PI3Kγ^−/−^ and PI3Kγ_inact_/PI3Kγ^−/−^ over WT. n = 5 to 6 mice per genotype. *P < 0.05 compared to WT and PI3Kγ_inact_/PI3Kγ^−/−^. **(C)** PLN immunoprecipitates from cardiac lysates of WT, PI3Kγ^−/−^ and PI3Kγ_inact_/PI3Kγ^−/−^ mice were immunoblotted for coimmunoprecipitating PP2Ac, PP1 and PI3Kγ and immunoblotted for PLN to show pull down of PLN by the antibody. **(D)** SR fractions of WT, PI3Kγ^−/−^ and PI3Kγ_inact_/PI3Kγ^−/−^ mice hearts were immunoblotted with anti-demethylated PP2Ac antibody. The blots were stripped and reblotted with anti-PP2Ac antibody. **(E)** Cumulative densitometric data is shown in the bar graphs. n = 5 to 6 mice per genotype. *p<0.05 compared to WT and PI3Kγ_inact_/PI3Kγ^−/−^. **(F)** SERCA2 immunoprecipitates from SR fractions of WT, PI3Kγ^−/−^ and PI3Kγ_inact_/PI3Kγ^−/−^ mice hearts were immunoblotted with anti–PLN antibody. The blots were stripped and reblotted with anti-SERCA2 antibody. **(G)** Amalgamated densitometry is shown in the bar graphs. n = 5 to 6 mice per genotype. *p<0.05 compared to WT and PI3Kγ_inact_/PI3Kγ^−/−^. **(H)** PLN immunoprecipitates from SR fractions of WT, PI3Kγ^−/−^ and PI3Kγ_inact_/PI3Kγ^−/−^ mice hearts were immunoblotted with anti-SERCA2 antibody. The blots were stripped and reblotted with anti–PLN antibody. **(I)** Amalgamated densitometry is shown in the bar graphs. n = 5 to 6 mice per genotype. *p<0.05 compared to WT and PI3Kγ_inact_/PI3Kγ^−/−^.

Since association of PLN with SERCA2 has consequences in calcium cycling, we tested if the unique regulation of PLN phosphorylation by PI3Kγ determines the assembly of PLN-SERCA2 complex. SERCA2 was immunoprecipitated and assessed for co-immunoprecipitating PLN. Significantly higher association of PLN was observed in the PI3Kγ^−/−^ mice compared to WT which was significantly reduced in the PI3Kγ_inact_/PI3Kγ^−/−^ mice (**Fig 5F & G**). Consistently, immunoprecipitation of PLN showed significantly higher association of SERCA2 in PI3Kγ^−/−^ mice compared to WT which was normalized in the PI3Kγ_inact_/PI3Kγ^−/−^ mice (**Fig 5H & I**). Together, these data show that PI3Kγ non-canonically alters calcium cycling in cardiomyocytes by regulating PLN phosphorylation through inhibition of protein phosphatases thereby, altering the assembly and function of SERCA2-PLN complex.

## DISCUSSION

This current study shows that PI3Kγ can regulate cardiac contraction by regulating the dynamics of calcium cycling and mobilization in the cardiomyocytes. Herein we have uncovered a pathway showing that PI3Kγ is an important player in calcium reuptake by SR. We further show that PI3Kγ aids in maintaining functional homeostasis of the SERCA2 via reuptake of calcium by regulating PLN through inhibition of the phosphatases. Thus, the mechanism of sustaining PLN phosphorylation by imparing phosphatase releases SERCA2 from PLN inhibition allowing for effective calcium cycling (**Fig 6**). Importantly, PLN phosphorylation is unexpectedly regulated by PI3Kγ through inhibition of phosphatase activity with minimal changes in phosphatase interaction with PLN. Intriguigingly, PI3Kγ mediates inhibition of PP2A through kinaseindependent mechanism as expression of kinase dead PI3Kγ_inact_ in the PI3Kγ null hearts (PI3Kγ^−/−^) normalizes phosphatase activity and restores of calcium cycling homeostasis. Consistently, the strong interaction observed between SERCA2 and PLN in the PI3Kγ^−/−^ hearts is reversed by expression of PI3Kγ_inact_ in the PI3Kγ^−/−^ hearts. Howerver, the association of PP1 or PP2A with PLN is not altered in the PI3Kγ^−/−^ hearts suggesting mechanisms beyond the kinase-independent scaffolding regulation of phosphatases by PI3Kγ (Mohan, et al. 2017). Finally, our studies show the presence of significantly low levels of demethylated PP2A in the SR of the PI3Kγ^−/−^ hearts (representing higher methylation mediated PP2A activity) which is reversed by expression of PI3Kγ_inact_ showing a previously unrecognized role of PI3Kγ in calcium cycling. This observation has significant implication in cardiac remodeling as PI3Kγ is upregulated in conditions of cardiac stress (Fougerat, et al. 2008; Patrucco, et al. 2004; Perino, et al. 2011; Perrino, et al. 2007).

**Fig 6.**
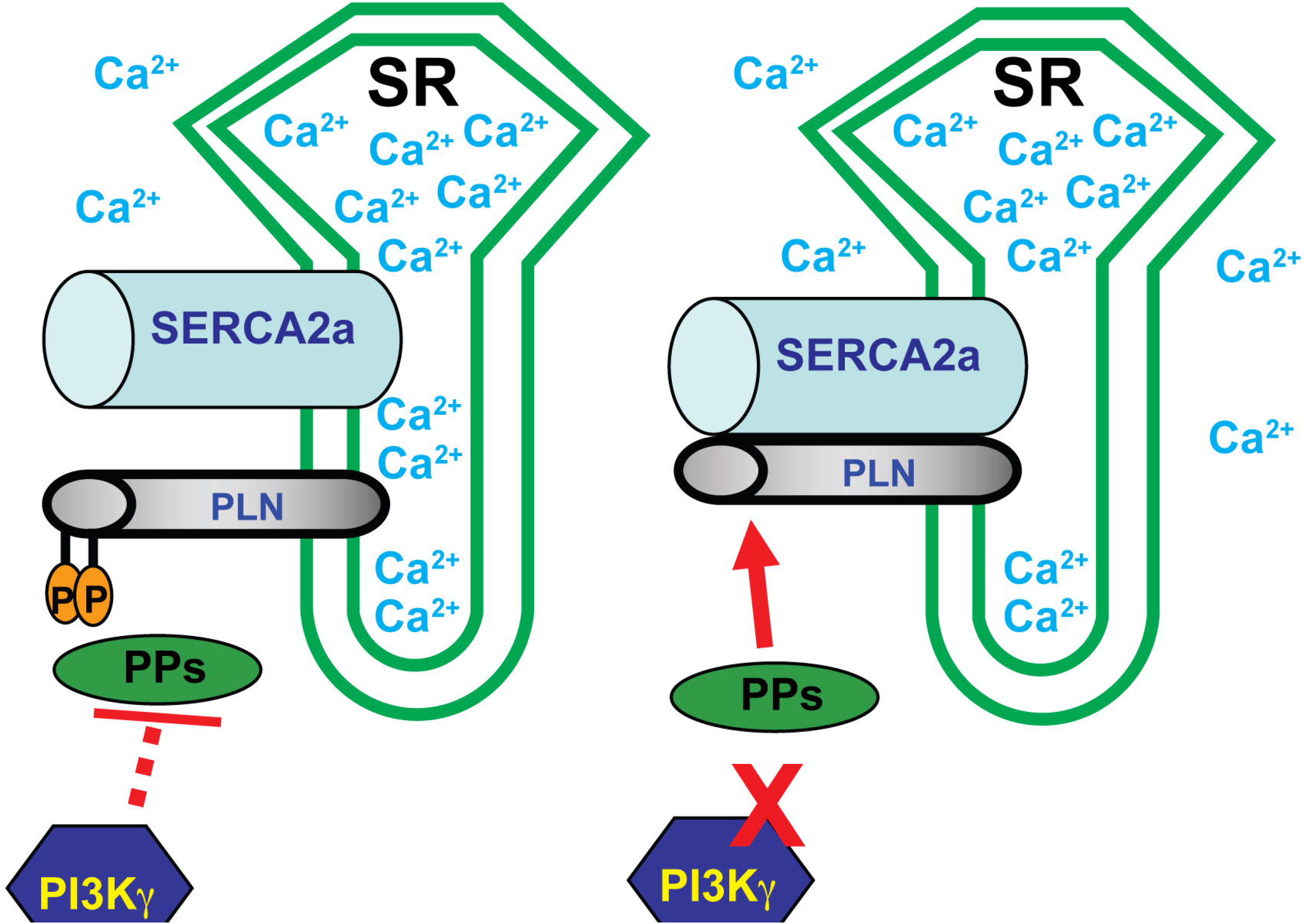
Graphical representation of PI3Kγ mediated regulation of PLN. PI3Kγ through inhibition of protein phosphatases enhance the phosphorylation of PLN. This leads to the release of inhibition on SERCA2 resulting in normal reuptake of calcium by SR. The absence of PI3Kγ leads to overactivation of protein phosphatases resulting in inhibition of SERCA2 culminating in resistance to normal reuptake of calcium by SR.

The molecular mechanisms regulating SR calcium transport and reuptake have been the subject of intense investigation as they are key regulators of cardiac contraction. Consistently, calcium mishandling is one of the characteristic features of failing myocardium (Bers, et al. 2003; Hasenfuss and Pieske 2002). The cardiac SR acts as a dynamic reservoir for calcium release and active uptake underlying cardiac excitationcontraction coupling and subsequent relaxation (Periasamy, et al. 2008). Our studies show that PI3Kγ plays an important role in regulating calicium cycling homeostasis. Previous studies have shown that absence of PI3Kγ leads to increased contractility (Alloatti, et al. 2005; Crackower, et al. 2002; Kerfant, et al. 2005; Nienaber, et al. 2003; Patrucco, et al. 2004) but, the underlying mechanisms have remained elusive. In this context, the observation of sustained intracellular calcium in the PI3Kγ^−/−^ cardiomyocytes outlines a unique yet to be understood integral role for PI3Kγ^−/−^ in this process.

Although multiple players have been recognized to play a role in calicium cycling homeostasis, SR RyR2 and SERCA pump pivotally contribute to the calcium release and reuptake. Previous studies have shown the RyR2 phosphorylation and dephosphorylation plays a critical role in regulating RyR2 dependent calcium release from SR (Ai, et al. 2005; Bovo, et al. 2017). In this regard, our study shows that RyR2 phosphoryation is not altered in the PI3Kγ^−/−^ cardiomyocytes. Correspondingly, caffeine treatment leads to high fidelilty release of calcum from the SR reserves showing that RyR2 function is not altered in the PI3Kγ^−/−^ cardiomyocytes. However, PI3Kγ^−/−^ cardiomyocytes are characterized by slow reuptake of calcium by the SR suggesting a role for SERCA2. Previous studies have shown that SERCA2 is functionally downstream of PI3Kγ in pancreatic acinar cells (Fischer, et al. 2007) and is also regulated by phosphorylation (Narayanan and Xu 1997; Xu, et al. 1997). Similar to RyR2, there was no appreciable difference in SERCA2 phosphorylation in the PI3Kγ^−/−^ cardiomyocytes elucidating that PI3Kγ does not directly regulate RyR2 or SERCA2 function. Interestingly, our study reveals that PI3Kγ alters the SERCA2 function through regulation of PLN as there is significant loss in PLN phosphorylation in the PI3Kγ^−/−^ cardiomyocytes.

PLN phosphorylation and dephosphorylaton is known to dynamically regulate SERCA2 function in calcium reuptake(MacLennan, et al. 2003; MacLennan and Kranias 2003). Abrogation of PLN phosphorylation in the PI3Kγ^−/−^ cardiomyocytes shows that the loss in the steady-state phosphorylation of PLN leads to increased inhibiton of SERCA2. This inhibition is reflected by the slow reuptake of calcium by the SR in the PI3Kγ^−/−^ cardiomyocytes. Consistently, significant interaction was observed between SERCA2 and PLN in the PI3Kγ^−/−^ hearts compared to the controls but surprisingly, this interaction was completely normalized by the expression of kinase dead PI3Kγ_inact_ in the PI3Kγ^−/−^ hearts. Corresponding to the loss in PLN phosphorylation, PI3Kγ^−/−^ hearts were characterized by augmented PLN-associated phosphatase activity which was unexpectedly reversed by PI3Kγ_inact_ in the PI3Kγ^−/−^ hearts. Moreover, inhibition of phosphatase by okadaic acid reverses Ca^2+^ reuptake by SERCA2 in PI3Kγ^−/−^ cardiomyocytes further supporting the idea that increased phosphatase activity is key underlying mechanism for dysregulated Ca^2+^ reuptake in PI3Kγ^−/−^ cardiomyocytes. These data show that PI3Kγ regulates SERCA2 function through PLN by inhibiting phosphatase activity by a kinase-independent mechanisms as phosphatase activity and calcium cycling are normalized by the expression of PI3Kγ_inact_ in the PI3Kγ^−/−^ hearts.

Although our studies reveal a unique kinase-independent role of PI3Kγ in regulating state of PLN phosphorylation, there was no difference in the association of phosphatases (PP1 or PP2A) with PLN in the PI3Kγ^−/−^ or PI3Kγ_inact_ in the PI3Kγ^−/−^ hearts. While PP1 is a well known regulator of PLN function, less is known about PP2A in regulating PLN dephosphorylation. In this regard, our previous studies have shown that PI3Kγ regulates PP2A activity and as absence of PI3Kγ (PI3Kγ^−/−^) leads slow reuptake of calcium indicates that PP2A also plays a role in PLN regulation. PP2A activity is well known to be regulated by methylation and demethylation, wherein methylation increases PP2A activity (Stanevich, et al. 2011; Xing, et al. 2008). Consistently, PP2A was significantly demethylated in the PI3Kγ^−/−^ hearts compared to wild type or PI3Kγ_inact_ in the PI3Kγ^−/−^ hearts. Low demethylated PP2A indicates higher methylation mediated PP2A activity. This suggests a yet to be appreciated role of PI3Kγ in regulating PLNassociated PP2A activity in SR through modulation of PP2A methylation instead of the kinase-independent scaffolding of PP2A. Thus, our study establishes that PI3Kγ maintains PLN function by impairing dephosphorylation of PLN by PP2A activity showing that PP2A function is integral to the homeostasis of calcium cycling in the cardiomyocytes. This homeostasis is critically driven by PI3Kγ as its absence (PI3Kγ^−/−^) leads to slow reupake of calcium, slow relaxation of cardiomyocytes and may potentially underlie a phenotype of diastolic dysfunction. These findings have implications in cardiac physiology because PI3Kγ expression is low in the normal healthy cardiomyocytes and is dramatically upregulated following stress suggesting another layer of PLN regulation by PI3Kγ-PP2A axis in maintenance of calcium homeostasis and cardiac contractile function.

## Acknowledgements

The studies are supported in part by AHA TPA (18TPA34170554) to MLM, R01 HL133721 and R01 HL 089473 to SVNP.

