## Supplementary material for "Kinase independent function of PI3Kγ modulates calcium re-uptake by regulating phospholamban function": Fig S1

**SUPPLEMENTAL MATERIALS**

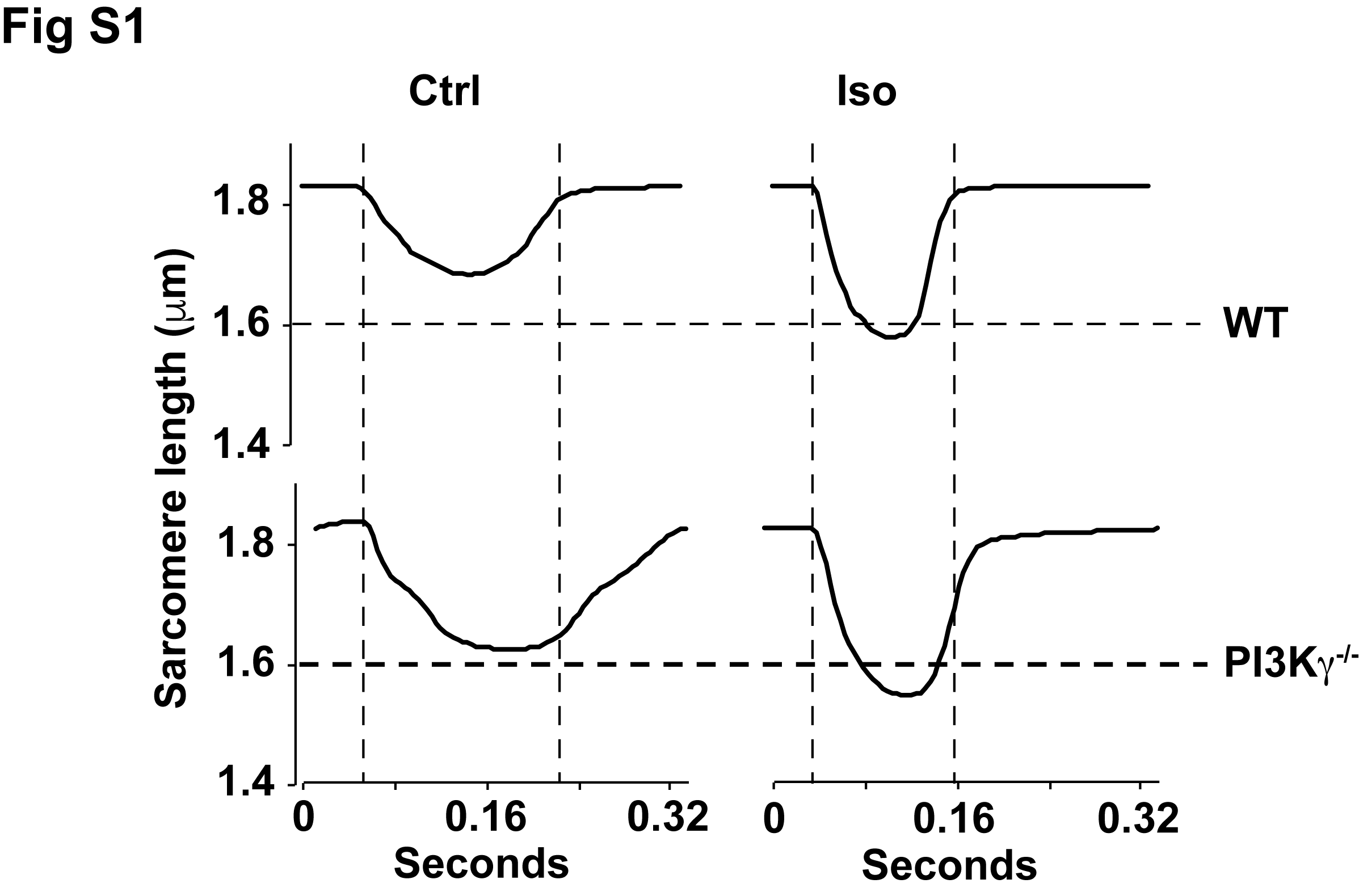

**Fig S1. Basal contractility is altered in isolated myocytes of PI3Kγ^-/-^ mice.** Average of cardiomyocyte contractility traces of WT and PI3Kγ^-/-^ cardiomyocytes (n=3 mice; 10-15 myocytes per mouse; 90-100 contractions/myocyte)
